# A deep learning framework for improving protein interaction prediction using sequence properties

**DOI:** 10.1101/843755

**Authors:** Yi Guo, Xiang Chen

## Abstract

**Motivation:** Almost all critical functions and processes in cells are sustained by the cellular networks of protein-protein interactions (PPIs), understanding these is therefore crucial in the investigation of biological systems. Despite all past efforts, we still lack high-quality PPI data for constructing the networks, which makes it challenging to study the functions of association of proteins. High-throughput experimental techniques have produced abundant data for systematically studying the cellular networks of a biological system and the development of computational method for PPI identification.

**Results:** We have developed a deep learning-based framework, named iPPI, for accurately predicting PPI on a proteome-wide scale depended only on sequence information. iPPI integrates the amino acid properties and compositions of protein sequence into a unified prediction framework using a hybrid deep neural network. Extensive tests demonstrated that iPPI can greatly outperform the state-of-the-art prediction methods in identifying PPIs. In addition, the iPPI prediction score can be related to the strength of protein-protein binding affinity and further showed the biological relevance of our deep learning framework to identify PPIs.

**Availability and Implementation:** iPPI is available as an open-source software and can be downloaded from https://github.com/model-lab/deeplearning.ppi

## INTRODUCTION

Interactions among proteins sustain largely critical biological processes happening in the cells (Westermarck, et al., 2013). Perturbations in those interaction networks can lead to disease (Sahni, et al., 2015). Determining accurate cellular networks of protein-protein interactions (PPIs) are therefore crucial for a proper understanding of various biological processes in cells for disease research and for drug discovery, as most common targets of drugs are proteins (such as enzymes, ion channels, and receptors) (Petta, et al., 2016).

A variety of computational methods have been developed to predict PPIs (Ding and Kihara, 2018). Computational methods based on genomic or proteomic information, such as phylogenetic profiles, coevolution rules, subcellular localization, and domain signature, predict PPIs by accounting for the pattern of the presence or absence of a given gene or protein. However, the accuracy and reliability of most of these methods depend on the information marks of the proteins in PPIs, which limits the universal predictive capacities of these methods. The primary amino acid sequence remains the most complete type of data available for all proteins. Thus, there is a longstanding interest in using sequence-based methods to model and predict protein interactions (Hashemifar, et al., 2018). Several sequence-based methods have been developed to predict PPIs, such as SigProd (Martin, et al., 2005), AutoCorrelation (Guo, et al., 2008), Yang’s work (Yang, et al., 2010), Zhou’s work (Zhou, et al., 2011), PIPE2 (Pitre, et al., 2012), You’s work (You, et al., 2014), PPI-PK (Hamp and Rost, 2015), Wong’s work (Wong, et al., 2015), Sun’s work (Sun, et al., 2017), and DPPI (Hashemifar, et al., 2018). The main limitation of these sequence-based methods is only the amino acid compositions of a sequence were considered.

Proteins directly interact via their three-dimensional (3D) structures (Vangone and Bonvin, 2015). However, exploring interaction processes at atomic level for a pair of proteins demand not just the knowledge of the 3D structure of the associated molecular. A typically interaction processes are highly restricted to the contact ways given the vast number of possible orientations (Cui, et al., 2019). One reason is that the amino acids of proteins encodes the information required to not only reproducibly fold them into their specific shapes, but to also reliably negotiate their biophysical interactions. The systematic analysis of PPI interfaces has shown complementarity in shape and electrostatic properties of amino acids (Sheinerman, et al., 2000). More recently, evolutionary coupling has demonstrated that coordinated changes in amino acids across evolution in pairs of proteins often occurs at interaction interfaces (Vangone and Bonvin, 2015). Taken together, the compositions and properties of amino acids of primary sequence of a protein informs both its structure and subsequent capacity to interact with other proteins.

Recently, deep learning continues to grow in popularity in computational biology and bioinformatics and has yielded the state-of-the-art performance over conventional machine learning on various types of biological prediction tasks, such as the predictions of protein-nucleotide binding (Li, et al., 2017), translation initiation sites (Zhang, et al., 2017), protein structure (Hou, et al., 2018), and functional effects of non-coding sequence variants (Zhou and Troyanskaya, 2015). In this study, we have developed a deep learning-based framework, named iPPI (identification of protein-protein interaction), for accurately predicting direct physical PPIs based on all protein-available primary sequences. iPPI possesses more reliability than previous methods, and integrates the composition of amino acids as well as the properties of amino acids in the primary protein sequences into a unified framework, in which an ensemble of hybrid deep neural networks (fully connected recurrent neural network) is implemented to effectively and robustly capture the sequence features of PPIs. Extensive validation tests have shown that iPPI can greatly outperform the state-of-the-art sequence-based approaches in detecting PPIs. By combining structural dynamics and energy data of mutated protein interactions with iPPI analysis, we found that the predictions of iPPI well correlated with the experimentally verified binding affinity of the protein-protein complexes, which basically demonstrated that iPPI can offer a powerful tool to model the direct physical contact between proteins and identify potential PPIs, which will improve the accuracy of determining cellular networks.

## METHOD

### Overview

In this study, we have developed a deep learning-based framework, named iPPI (identifying protein-protein interaction), for accurately predicting PPI based on the properties of amino acids on the protein primary sequences. Figure 1 illustrates the schematic overview of our deep learning framework, which consists of three main phases, including problem statements about feature and dataset, design solutions for the problem, and training models to improve prediction. In the problem statement phase, we first propose the amino acid properties of PPI closely related protein sequences and the influence of imbalanced training set on prediction accuracy. Then, we encode the primary sequence with the properties of 20 amino acids and employ a random undersampling strategy to solve the problems. In the training phase, we build a hybrid deep neural network to model the encoded sequence profiles of the bootstrapping-based training datasets.

**Figure 1.**
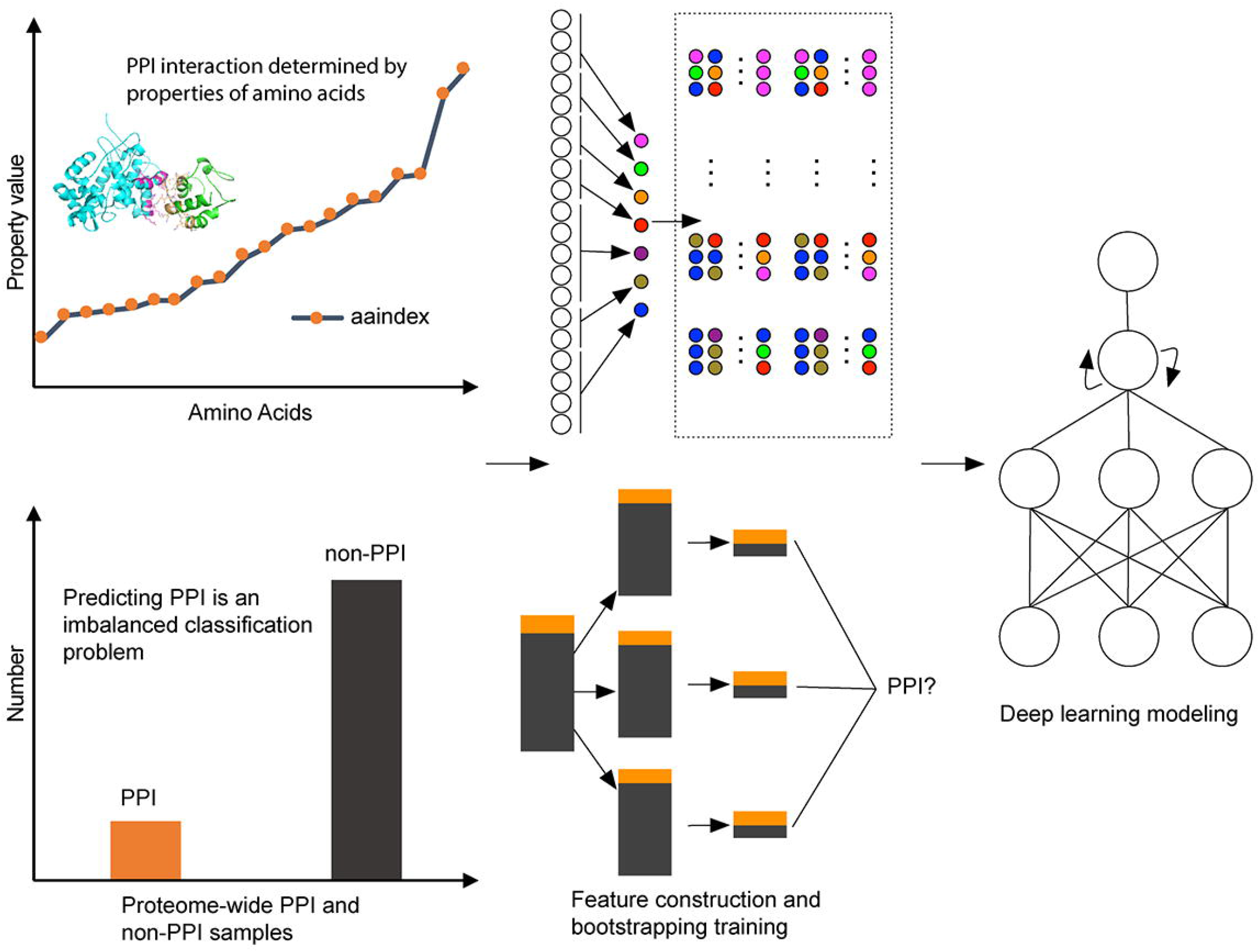
Schematic overview of the iPPI pipeline. See the main text for more details

### Encoding PPI pair sequences

For predicting PPI by sequences, one of the main computational challenges is to find a suitable way to fully describe the important feature information of PPI. To solve this problem, we proposed a descriptor named conjoint aaindex modules (CAM), which considered any conjoint amino acids (CAA) of protein sequences as a unit and encoded the unit with AAindex database that defines the properties of amino acids. Thus, the CAA can be categorized as CAM according to the classification of the property value, i.e., For different conjoint amino acids “CAC” and “CCC”, if “C” and “A” are assigned to the same group by the property value, then “CAC” and “CCC” are grouped together and considered to be the same unit for protein sequence. For a given primary protein sequence with length n we generate a homogeneous vector space by counting the frequencies of each CAM type appearing in the protein sequence, called sequence profile, as follows:

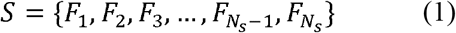

where *S* is the frequency vector that represents the frequencies of CAM type in the protein sequence, and *N_s_* is the number of CAM type. The process of generating the vectors is described as follows. First, the amino acids were catalogued into assigned sizes of clusters by using *k*-means method to cluster the property value. Each amino acid will be labeled based on this classification. Second, the protein sequences were replaced with the labels to generate label sequences. Third, a sliding window of fixed length was selected to find patterns of CAA, also known as CAM. The number of CAM types varies depending on the window length. Finally, the frequencies of each CAM type were computed to form the frequency vector *S* for the given protein, as follows:

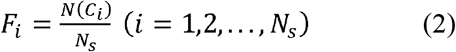

where *C*_*i*_ represents the CAM type and *N*(*C*_*i*_) is the number of the CAM type. Figure 2 illustrates the calculation process by taking the amino acids divided into seven clusters and the window length set as three. In this result, the size of vector *F* should be 343 (=7*7*7). Long protein sequences usually have larger frequency values than short protein sequences, which may adversely affect the performance of learning algorithms. To solve this problem, min-max normalization procedures are used that ensures the *F_i_* are between 0 and 1. iPPI takes a pair of protein profiles *S*_*A*_ and *S*_*B*_. Considering the fact that PPI is symmetrical, i.e., {*S*_*A*_,*S*_*B*_} and {*S*_*A*_,*S*_*B*_} represent the same interaction pairs between proteins A and B, we designed a function to concatenate the vector spaces of two proteins to represent their interaction features: *K* = (*S*_*A*_ − *S*_*B*_)^2^. Thus, a protein pair vector of the same dimension as a protein vector was generated to represent each protein pair. Finally, this vector was used as the input fed into deep learning framework to output a binary value indicating whether the corresponding proteins interact.

**Figure 2.**
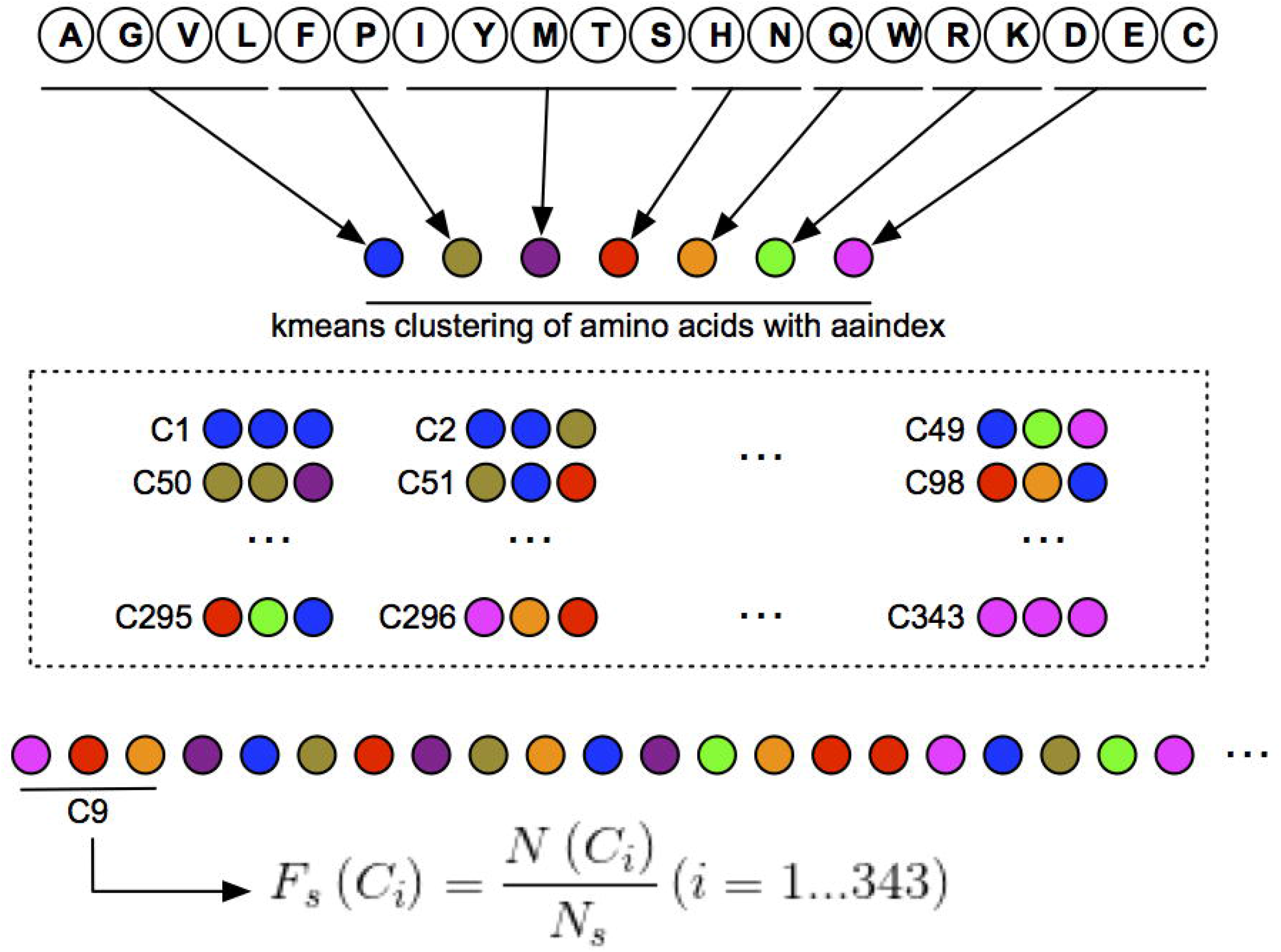
Schematic illustration of encoding protein primary sequence

Clustering amino acids properties: AAindex is a database of numerical indices representing various physicochemical and biochemical properties of amino acids, which currently contains 566 amino acid indices classified in six groups: alpha and turn propensities, beta propensity, composition, hydrophobicity, physicochemical properties, and other properties. Each of these amino acid properties derived from published literature is referred to as an entry. Each entry consists of an accession number and a set of 20 numerical values corresponding to 20 amino acids. As above mentioned, *k*-means is used to cluster the numerical values. To perform this, 20 numerical values are sorted from large to small forming a numerical interval, and the equidifferent values of the assigned cluster number of the clustering from the numerical interval are selected for the starting value of the clustering. If the number of the assigned cluster size is larger than the kinds of the property value of 20 amino acids, the clustering will fail that leads the property cannot be used to predict the model.

### Deep learning framework

Here, we develop a deep neural network to systematically model the sequence features of PPI (Figure 3A). To characterize the feature of the PPI, we first use the CAM descriptor to encode the primary sequences of the protein pair as mentioned above. Then we employ multiple dense operators (denoted by *dence*()) to scan the encoded sequence profile *S* and detect the CAM patterns around each PPI. After that, the rectified linear unit (*ReLU*()) (Diederich, 2007) takes the output of a dense layer and clamps all the negative values to zero to introduce non-linearity. Note that the conventional dense-layer structures are order insensitive, as they only detect whether certain CAM exists regardless of their positions or orders. To enable order dependence can be captured that may also contribute to PPI, LSTM (long short-term memory, denoted by *LSTM*()) network (Hochreiter and Schmidhuber, 1997) is stacked upon the dense layer, which takes feature vectors from the dense layer as input and learns the long short-term dependencies at the CAM. Finally, the outputs of the LSTM are fed into a logistic regression layer (denoted by *logist*()) (Dreiseitl and Ohno-Machado, 2002) to compute the probability of PPI for the input sequence profiles. Altogether, by fully exploiting both sequence profiles and CAM orders, the hybrid fully connected and recurrent neural network in iPPI computes the following score:

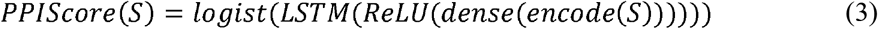

**Figure 3.**
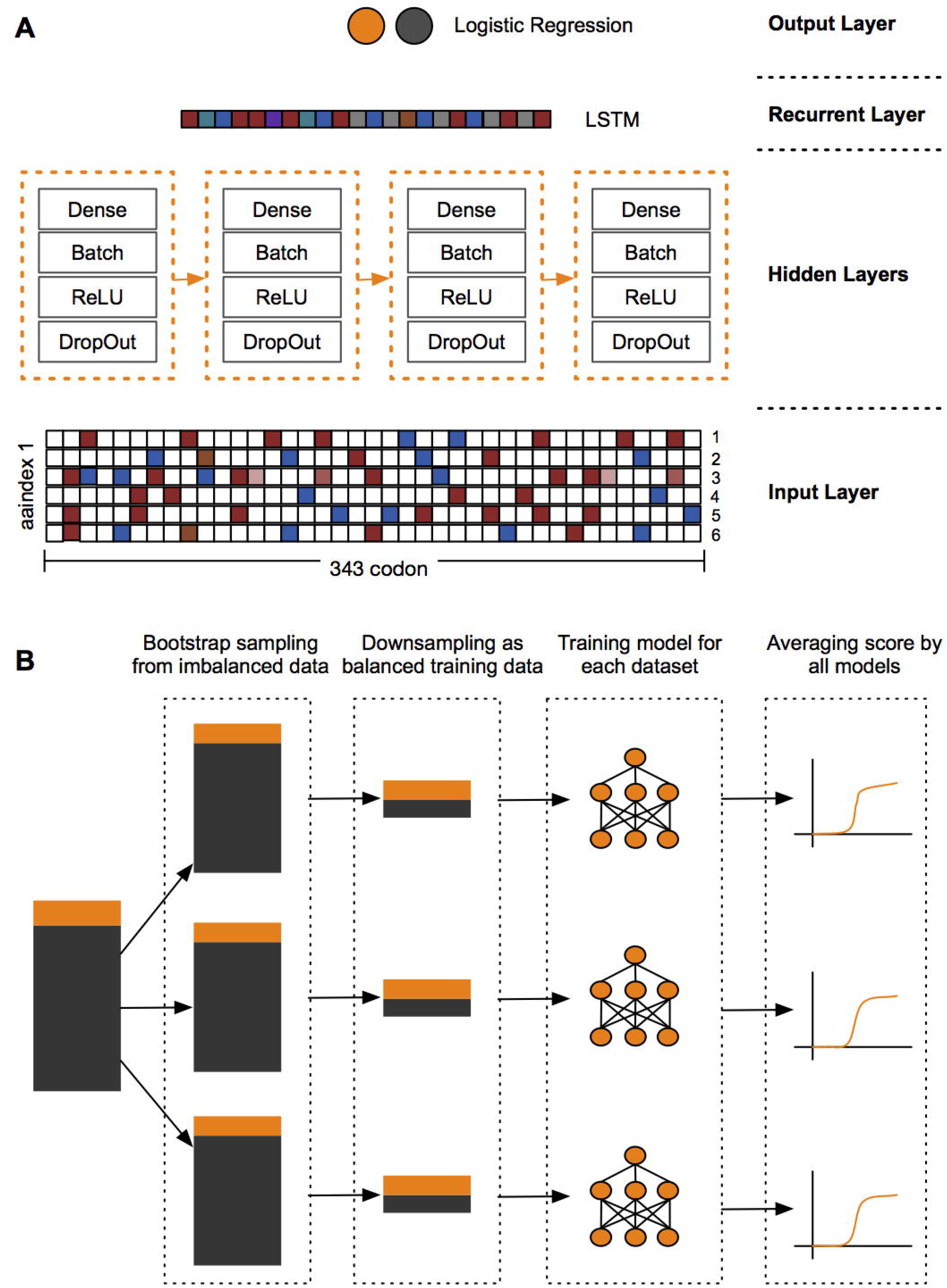
Schematic illustration of (A) the hybrid deep neural network architecture and (B) the bootstrapping-based technique used for imbalanced classification problem. See the main text for more details

The sequence features of non-PPI (i.e. the negative samples) can be modelled in the same manner. The details of the proposed framework are shown in Figure 3A. Note that a similar hybrid neural network architecture has also been implemented in DeepSEA (Zhou and Troyanskaya, 2015) and DeepBind (Alipanahi, et al., 2015) for different tasks, i.e. predicting the functional effects of noncoding variants and the sequence specificities of DNA- and RNA-binding proteins, respectively.

### Model training and model selection

Given the PPI training samples {((*S*_*A*_,*S*_*B*_), *y*_*i*_)}, where (*S*_*A*_,*S*_*B*_) is the input protein profiles, and *y*_*i*_ is the binary label that indicates whether is a pair of interacting proteins. The loss function *L*(*f*_*i*_,*y*_*i*_) of our deep learning framework is defined as the sum of the negative log likelihoods, i.e.

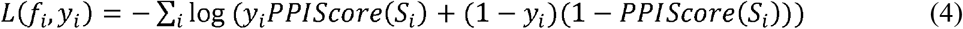

where *f*_*i*_ equals *f*_*i*_(*S*_*A*_,*S*_*B*_) that indicate the prediction of our model. We use the standard error backpropagation algorithm (Rumelhart, et al., 1986) and the stochastic gradient descent method (Kingma and Ba, 2015) to train the hybrid neural network and search for the network weights that minimize the loss function of Equation (4). The regularization techniques, dropout method (Srivastava, et al., 2014) are also implemented for dense layer to address the overfitting problem that regards the generalization capacity for predicting the unknown dataset (independent samples that are not in the training samples).

Our hybrid deep neural network architecture and the optimization techniques used in the training process have introduced several hyperparameters, e.g. the learning rate, layer size, and regularization constant, that need to be determined. A proper hyperparameter calibration procedure can help yield better solutions to the optimization problem in Equation (4). Here, we use the random search (RS) approach (Bergstra and Bengio, 2012) to calibrate the hyperparameters in our model, including learning rate, layer size, regularization constant, output dimension of the layer, and dropout rate (Srivastava, et al., 2014) in the dense layer. In particular, we first use all the positive training samples and an equal number of randomly selected negative training samples for our collected dataset to optimize the hyperparameters based on RS (with 10,000 evaluations), and then choose the hyperparameters that achieve the minimum loss to further train our final models.

We realize that the task of predicting PPI is an imbalanced classification problem (i.e. much more negative samples (non-PPI) than positive ones), for which the standard training procedure designed for balanced samples cannot be applied directly. On the other hand, making use of more negative samples in the training process can lead to a more robust model with less variance in prediction (Wallace, et al., 2011). To tackle this imbalance problem, here we employ a bootstrapping-based technique (Figure 3B) derived from the theory established in Wallace et al. (Wallace, et al., 2011). Briefly, we first construct several groups of bootstrap samples (denoted by *G*_*i*_) from the original imbalanced population *G*, by randomly selecting samples with replacement. Then for each group *G*_*i*_, we balance the samples by downsampling, i.e. randomly selecting an equal number of positive and negative samples from *G*_*i*_, which yields a balanced dataset *B*_*i*_. After that, a hybrid deep neural network *N*_*i*_ is trained based on each dataset *B*_*i*_ independently, resulting in an ensemble of binary classifiers {*N*_*i*_}. Given input sequences *S*, its final prediction score is averaged over the prediction scores output by all classifiers, i.e.

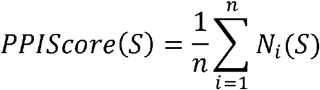

in which *n* is the total number of the constructed balanced datasets in *B*_*i*_ (which is also equivalent to the total number of trained classifiers in {*N*_*i*_}). We apply this bootstrapping-based technique to our training dataset and train 32 independent deep neural networks. After that, their prediction scores are averaged and used as the final estimated probability of translation initiation for the given input sequence profiles.

Owing to the nature of non-convex optimization, random weight initialization can influence the search results of the gradient descent algorithm (Kingma and Ba, 2015). This initialization bias may also introduce variance to our modelling and further affect the prediction performance. In iPPI, the aforementioned bootstrapping-based technique can alleviate such an initialization bias in addition to the sample bias, as the network weights have been initialized independently for each balanced sample group before the training process.

The hybrid deep neural network of iPPI has been implemented using the dl4j package (https://deeplearning4j.org).

## RESULTS

### Construction of predictors

We first optimized the cluster size for the clustering of the property values of 20 amino acids by traversing the sizes from five to nine and the length of the sliding window for find patterns of CAA by traversing the window size from three to five. In this work, PPI data from the Human Protein References Database (HPRD, version_20181103, www.hprd.org) (Peri, et al., 2004) that contains 37,961 PPI pairs as positive samples, which merged with an equal number of randomly selected negative samples as training set. The training set was encoded by each of cluster-available entries in AAindex to search the optimal results using a fully-connected deep neural network with three hidden layers. As shown in Figure S1, the cluster size and the sliding window length has brought a tremendous influence on the accuracy of PPI prediction. The optimal window size and cluster size are three and seven respectively, which are used in subsequent model optimization and analysis. Accordingly, 495 cluster-available entries were obtained; a PPI pair could be encoded into a 343-dimensional vector for a cluster-available entry in AAindex. To capture optimal entries for predicting PPI, the training set was encoded by each of the 495 cluster-available entries, and the fully-connected deep neural network as mentioned above was used to construct models for PPI predictions. The parameters of the network were set by the hyperparameter calibration procedure with 1,000 evaluations for each entry, respectively. The results are shown in Figure S2. The optimal entries for the six groups in AAindex were listed in Table S1 and Table S2, which show the amino acids of seven clusters and the clustering value of each cluster. To evaluate prediction ability in an unbiased manner, five independent datasets, from including Venkatesan et al. (Venkatesan, et al., 2009), Colland et al. (Colland, et al., 2004), Rual et al. (Rual, et al., 2005), Albers et al. (Albers, et al., 2005), and Bell et al. (Bell, et al., 2009), that were adopted as testing sets to retest these entries and their combination. As mentioned in the methods section, the hybrid deep neural network with a hyperparameter calibration procedure was designed for this test. The evaluation results have shown in Table S3 and Table S4, which have indicated a high prediction stability of PPI encoding by selected entries, and the optimal prediction performance performed on the combination of these selected entries. In addition, we also compared the prediction capacities of the combination with the CAM without the encoding of properties of amino acids (sliding window size was set three, each PPI pair was encoded as 20*20*20-dimensional vector), which show that the combination achieves a vast improvement (Table S5). Thus, the combination encodes each PPI pair into 343*6-dimensional vector that was chosen as the final encoding for the following model construction on PPI prediction. Based on the encoding above, we next sought to test the prediction performance for different size of network layers and different number of output neurons of each network layer in the hybrid deep neural network. The results are shown in Figure S3. First, we evaluated the prediction capacities of the different size of network layers through the hyperparameter calibration procedure with 6,000 evaluations. Interestingly, the number of network layers reaches the best prediction at seven layers, while the prediction performance weakens after seven layers (Figure S3A). Second, we further evaluated in more detail to obtain the optimal size of network layers using the network architecture from five to seven layers. The results were observed through the hyperparameter calibration procedure with 10,000 evaluations, which indicate that seven-layer network architecture has the optimal prediction capacities. Finally, the seven-layer network architecture was used to evaluate the number of output neurons for each network layer. The number of output neurons for each dense layer has been exhibited in Figure S3C that were introduced final deep learning framework to construct the predictor called iPPI for the detection of potential PPI. All network-related parameters in the framework are listed in Table S6.

### iPPI accurately predicts PPIs

To evaluate the prediction performance of iPPI, the *in vivo* human PPI data from Hippie database v2.2 (Alanis-Lobato, et al., 2017) was adopted as positive samples of training set, and five independent datasets as mentioned above excluded protein sequences present in the training set that was used as positive samples of testing sets. Overall, the training set has 18,738 PPIs, and the testing sets have 182 PPIs from Venkatesan et al. (Venkatesan, et al., 2009), 759 PPIs from Colland et al. (Colland, et al., 2004), 2,845 PPIs from Rual et al. (Rual, et al., 2005), 229 PPIs from Albers et al. (Albers, et al., 2005), and 2894 PPIs from Bell et al. (Bell, et al., 2009). As mentioned in the section of “Model training and model selection”, negative samples with the same number of positive samples were selected by the bootstrapping-based technique to construct training and testing sets. Based on this, we set each training set with 1,600 samples and constructed 32 training sets. In the training stage of iPPI, 32 models were trained and assembled to detect PPI based on the 32 training sets and the parameters in Table S6. After that, we performed 32 predictions for each of the five testing sets. Figure 4 presents the mean auROC (area under the ROC curve) and mean auPR (area under precision-recall curve) of the 32 predictions on the different testing sets. In summary, the results showed that iPPI achieved an acceptable performance for independent test data of different sizes. The auROC and auPR were mostly greater than 0.70. Importantly, the sensitivity reached 0.30-0.56 when the false positive rate was 0.1, and the precision was 0.53-0.65 when the recall (predictive sensitivity) was set to 0.9. Interestingly, we found that test sets with larger data sizes, such as dataset from Colland et al., achieved better predictive results, which further indicate that the deep learning framework of iPPI is accurate and robust in predicting PPI. Finally, we performed extensive tests on a variety of datasets to compare with other methods (supplemental information for details). The results show that iPPI outperformed the state-of-the-art sequence-based methods (Figure 5, Figure SS1, and Table SS1).

**Figure 4.**
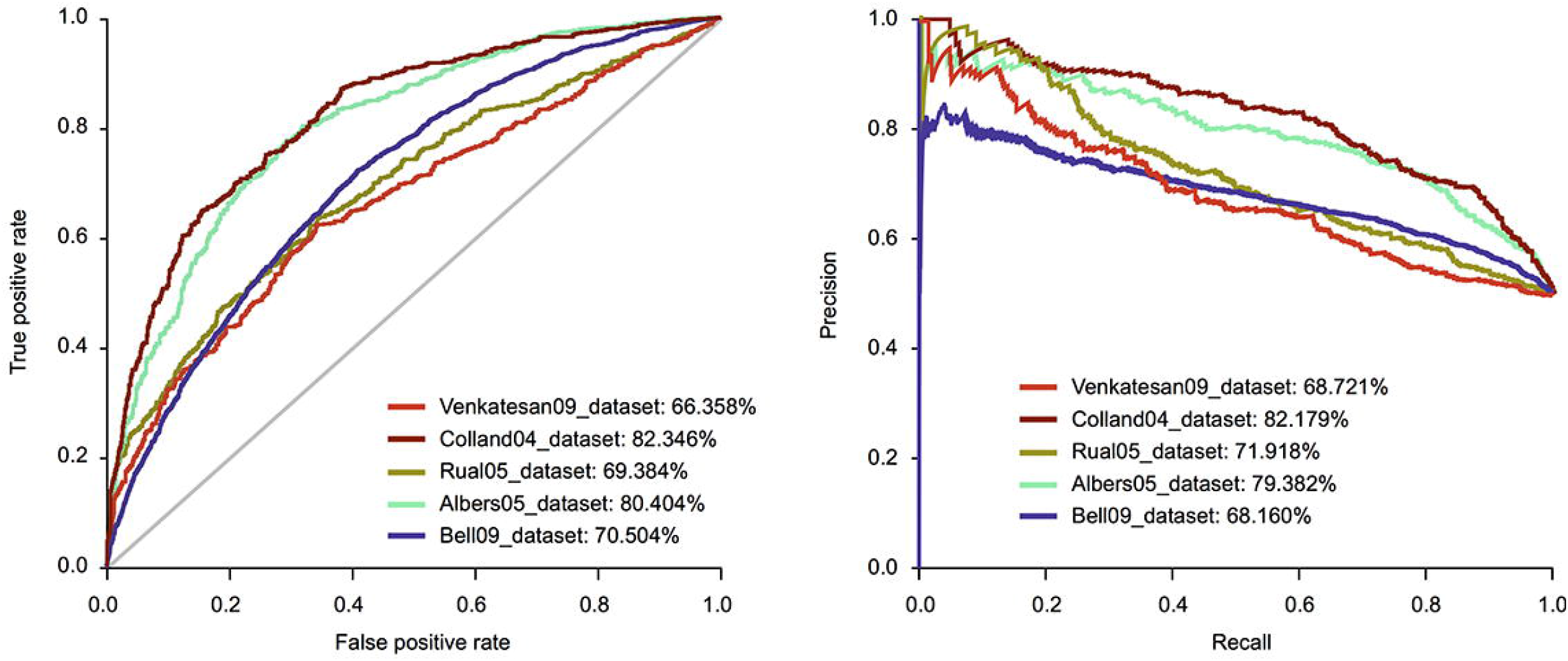
Prediction performance on different test datasets. The prediction performance on the different datasets evaluated by (A) ROC and (B) PR curves. The digits in brackets represent the proportion of the area under the curve

**Figure 5.**
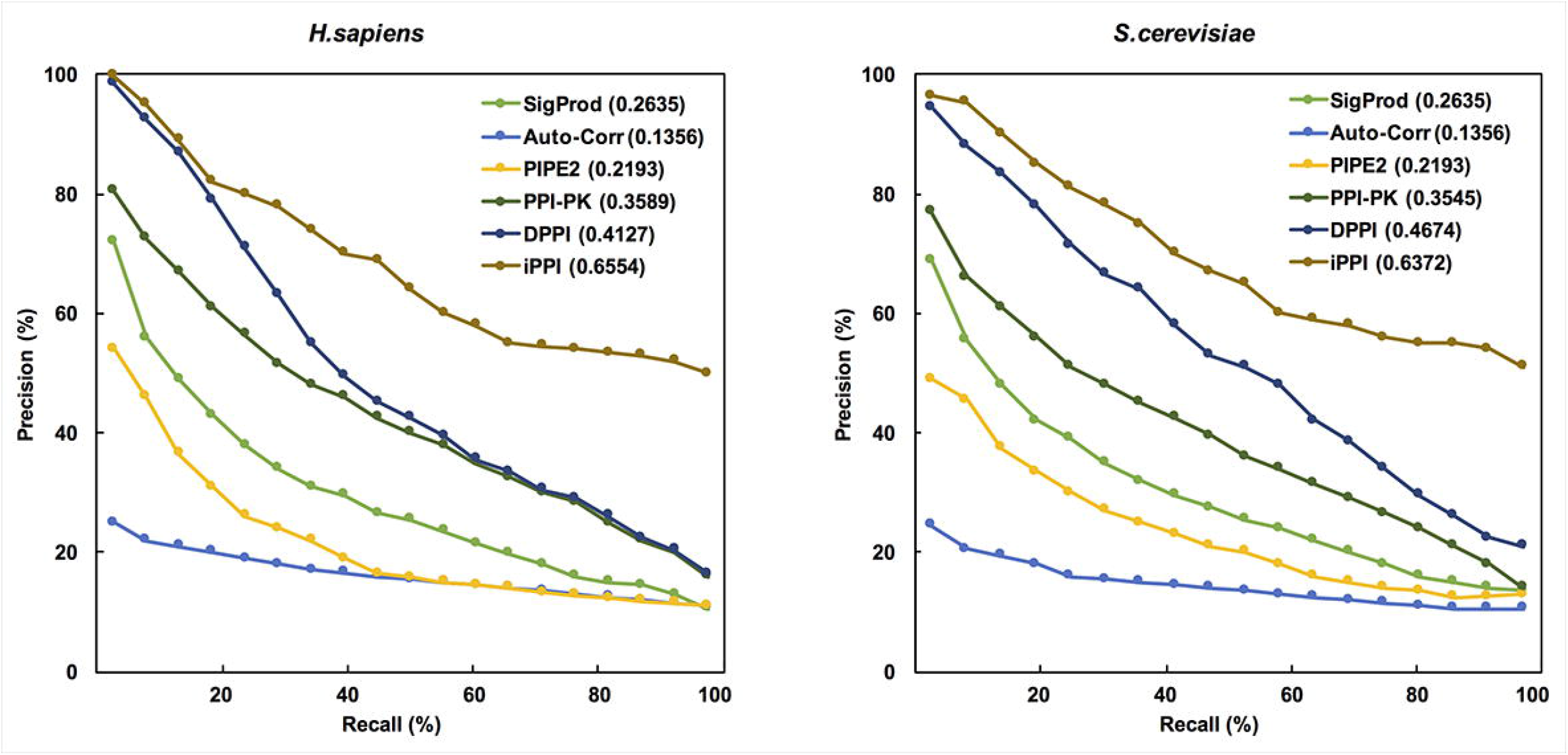
The comparison of the prediction capacity of different tools on the test datasets of both *H. sapiens* and *S. cerevisiae* using 10-fold cross-validations

### CAM-based prediction of iPPI correlates with binding affinities

As introduced in the method section, we obtained 495 cluster-available entries when optimizing the number of cluster categories, which belong to six different groups delimited by AAindex. According to the observation of Figure S4, the capacity of different groups of entries to predict PPIs has large differences. For the median, the group of physicochemical properties obtained the best prediction performance, and the worst performance presented in the group of beta propensity. However, the group of hydrophobicity category performed best in terms of prediction accuracy. As above mentioned, we finally selected the six best entries from these different groups in order to fully capture the background of PPIs. The entries we use include JANJ780101, MIYS990102, MIYS990102, MIYS990102, RICJ880109, OOBM770104 that respectively belong to hydrophobicity, other properties, beta propensity, composition, alpha and turn propensities, and physicochemical properties. We further analyzed the differences between positive and negative samples of these entries encoded CAM (Figure S5). This result combined with Table S1 and S2 provides some interesting observations that the enrichment patterns of CAM showed differences in positive and negative samples. For example, the patterns of “315” and “133” are shown in positive samples, while a symmetric pattern “311” and “113” are concentrated in negative samples for the entry of JANJ780101. More interestingly, both cluster 3 and cluster 1 appear frequently in the enriched CAMs for the positive and negative samples. The JANJ780101 describes the average accessible surface area of amino acids, which plays a key role in the conformation of protein side chains (Janin and Wodak, 1978). The side chain of protein determines the interface of protein-protein contact has been confirmed in many studies (Geppert, et al., 2011; Xue, et al., 2015), thus we supposed that this is why the CAM patterns derived from the JANJ780101 can effectively predict PPIs.

It has been indicated that the conservation of binding residues can contribute strongly to binding energetics (Brender and Zhang, 2015). Therefore, the properties of the residues (amino acids) play an important role in many prediction programs which seek to identify binding affinity and binding site on the structure interface of protein (Moreira, et al., 2007; Vangone and Bonvin, 2015). Since the encoding CAM used by iPPI considers the different amino acid properties of the proteins in PPIs, it may naturally reflect the direct physical contact between the two proteins. Therefore, we considered whether our model could effectively predict affinity between proteins. In this work, we defined the effect of mutations on the proteins of PPIs as the Gibbs free energy change (ΔΔ*G*) of binding between proteins. Quantitatively, ΔΔ*G* values for PPI may be measured experimentally by a variety of biophysical method (Kastritis and Bonvin, 2013). We first employed the 32 models defined in the section of “iPPI accurately predicts PPIs” (implemented based on the mean probability values of 32 models) to predict the probability values of PPI for the primary sequences and the mutation sequences on the PPI proteins. We sought to evaluate the correlation between the probability difference of the two predictions and the experimentally derived ΔΔ*G* values. Experimental ΔΔ*G* values were derived from the SKEMPI database that consists of experimental protein affinity changes upon mutation for protein-protein complexes in which a crystal structure of the WT complex is available (Moal and Fernandez-Recio, 2012). Following Brender et al. (Brender and Zhang, 2015), the redundant data was removed to obtain the final dataset containing 1,725 entries for 130 complexes. Test results showed that the probability difference based on the predicted probability values of PPIs computed by iPPI well correlated with the experimentally measured ΔΔ*G* values (the Pearson’ s correlation coefficient 0.604-0.683 (mean value: 0.646); Figure 6A), indicating that iPPI can provide predictive power to estimate the strength of binding affinities. Interestingly, we observed that the probability score of protein-protein complexes was significantly higher (*p-*value=5.2e−5, two independent sample t-test computed by comparing the distribution of probability scores) than the PPIs in the *in vivo* human PPI data from Hippie database v2.2 (Alanis-Lobato, et al., 2017) (Figure 6B). As expected, the results maintained a good consistency with the fact that the binding affinity of the protein-protein complexes is greater than the common physical contact, which thus further validated the biological relevance of our deep learning framework to identify PPIs.

**Figure 6.**
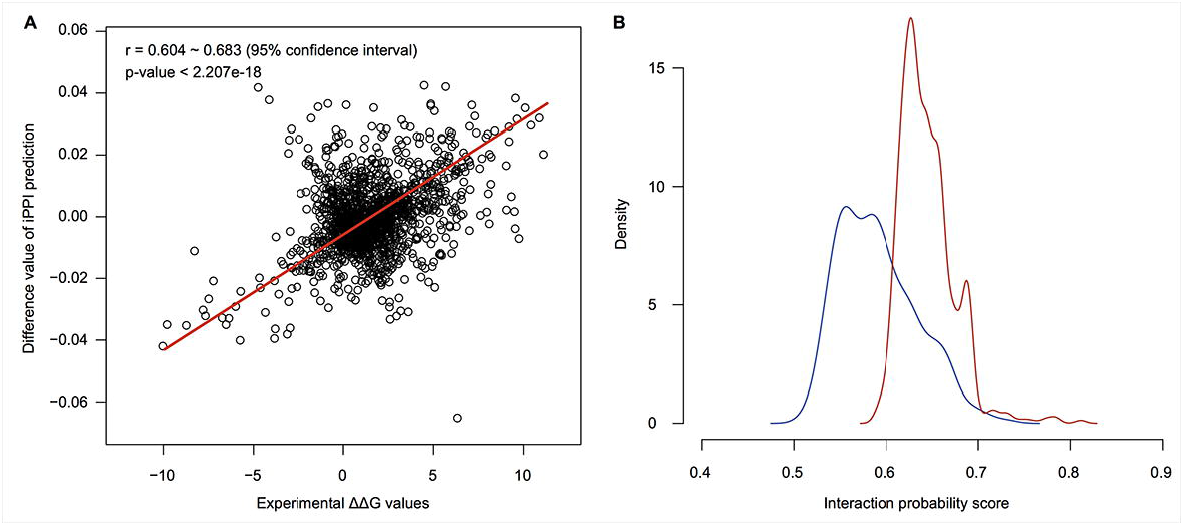
The probability scores of PPIs on protein-protein complexes. (A) The prediction of iPPI correlates with the experimentally measured mutational effects. (B) The comparison of the predicted probability scores between all PPIs from the *in vivo* human PPI data in Hippie database v1.2 (blue line) and protein-protein complexes from the SKEMPI database (red line)

## CONCLUSION

We have developed a deep learning framework to model the CAM preferences of interaction proteins by encoding the primary sequence with the properties of amino acids. Our framework considered six best entries from six grouped properties in AAindex including hydrophobicity, other properties, beta propensity, composition, alpha and turn propensities, and physicochemical properties, which fully capture the sequence features of PPIs. Tests on real independent datasets showed that our framework can achieve the comparable or superior performance to the state-of-the-art method for predicting PPIs. In addition, the encoding CAM considered CAA of primary sequences that means the information of non-synonymous mutation can be captured for PPI, which is applicable to different biological problems, such as binding affinity changes upon mutation for protein-protein complexes. The results also show that the binding affinity changes generated by our framework agreed well with the previous experimental studies, and may provide useful hints for further elucidating the molecular mechanisms of PPIs. In summary, our deep learning framework serves as a principled computational approach to model and predict PPIs from sequence and is generalizable to many applications.

## Supplementary figure legends

**Figure S1.** The accuracy of prediction by traversing the cluster sizes from 5 to 9 and the sliding window sizes from 3 to 5 (represent by different colors from left to right). Each of cluster-available entries in AAindex was used to encode train dataset for each traversal

**Figure S2.** The sorted accuracy of prediction by setting the window length as 3 and the cluster size as 7. By clustering properties, the train dataset encoded by each of 495 cluster-available entries in AAindex was used in this analysis

**Figure S3.** The optimization processes of model for the size of network layers and the number of output neurons of each dense layer. (A) the distribution of the frequencies of model score >50% for the different size of network layers. (B) the scores (prediction accuracy) on the top three layer sizes of the frequencies. (C) the optimization of the number of output neurons for six dense layers of seven network layers. The interval of each dense layer was set from the optimization in the step (B). The hyperparameter calibration procedures were used in these optimization processes

**Figure S4.** The comparison of prediction performance on other species datasets

**Figure S5.** The comparison of prediction capacities of the entries on different groups of property. The boxplot shows the predicted accuracy distribution of entries grouped in different categories. The prediction accuracy of all entries is derived from the results of Figure S1

**Figure S6.** The differences between positive and negative samples of CAM based on the encodings of the best 6 entries. The in vivo human PPI data from Hippie database v1.2 (Schaefer et al., 2012) mentioned in the main text was used to extract CAM. Each three-digit number in the figure represents a CAM, and each number in the three-digit number represents the cluster ID corresponding to the entry. The enrichment value greater than zero indicates a positive sample, and vice versa

## Supplementary table legends

**Table S1.** The amino acids of each cluster for the top six entries of prediction accuracy in each group of AAindex. The sorted accuracy of prediction was obtained from Figure S2. The entries include JANJ780101 (Janin and Wodak, 1978), MIYS990102 (Miyazawa and Jernigan, 1999), NAGK730102 (Nagano, 1973), RICJ880109 (Richardson and Richardson, 1988), OOBM770104 (Oobatake and Ooi, 1977)

**Table S2.** The clustering values for the top six entries that associate with the clusters in Table S1

**Table S3.** the prediction results of each of selected entries in AAindex and their combination that implement by the hybrid deep neural network framework mentioned the manuscript (the parameters of the network were set by the hyperparameter calibration procedure with 1,000 evaluations, respectively). These results were computed by 10-fold cross validations

**Table S4.** The *p*-values were computed by the comparisons of prediction accuracy for selected six entries in AAindex and their combination. The data associate with Table S3

**Table S5.** The comparison of prediction capacities between CAM with six entries and CAM with 20 amino acids

**Table S6.** The parameters of the deep learning framework of iPPI

**Table S7.** Performance comparison of iPPI with other state-of-the-art methods on the *S. cerevisiae* core subset. Other tools reported results from Hashemifar et al., 2018. Note: Bold font is used to indicate the best performance

