## supplemental information for "A deep learning framework for improving protein interaction prediction using sequence properties"

**(Supporting information)**

**PPI prediction quality** **comparing with other methods**

Here, we performed extensive tests on a variety of datasets to show that iPPI greatly outperformed the state-of-the-art sequence-based methods in predicting PPI, including SigProd (Martin, et al., 2005), AutoCorrelation (Guo, et al., 2008), Yang’s work (Yang, et al., 2010), Zhou’s work (Zhou, et al., 2011), PIPE2 (Pitre, et al., 2012), You’s work (You, et al., 2013), You’s work (You, et al., 2014), PPI-PK (Hamp and Rost, 2015), Wong’s work (Wong, et al., 2015), Huang’s work (Huang, et al., 2015), and DPPI (Hashemifar, et al., 2018). Note that some sequence-based methods were not considered for comparison because neither of their methods nor their data is currently available. In this work, we referred to DPPI and used *H. sapiens* PPIs obtained by collecting the 10% top-scoring interactions from Hippie database v1.2 (Schaefer, et al., 2012) as well as *S. cerevisiae* PPIs extracted from the DIP core set (Salwinski, et al., 2004) as test datasets. The redundant data with sequence identity greater than 0.4 for any protein in both PPIs were removed from the test datasets using the cd-hit tool (Fu, et al., 2012). Following the strategy used in DPPI (Hashemifar, et al., 2018), the same number of negative samples were selected 10 times to match positive samples as 10 training sets by randomly sampling from all proteins. We first tested our method using a 10-fold cross-validation method for each of 10 training sets and evaluated its prediction performance based on the mean auPR. The same test we used that is performed for other methods that had been reported in DPPI (Hashemifar, et al., 2018). To ensure that the comparison was unbiased, all results for these sequence-based methods were used. As presented in Figure 5, our model yielded better prediction performance (achieved the highest auPR value) on the test datasets of both *H. sapiens* and *S. cerevisiae*. We observed that our method greatly increased in auPR values by 0.2427 for *H. sapiens* and 0.1698 for *S. cerevisiae*, respectively. This indicated that the sequence-based CAM features using the properties of amino acids may provide more predictive information in our deep learning framework than other prediction tools.

We also performed additional tests on an independent dataset described by You et al. (You, et al., 2014) and DPPI (Hashemifar, et al., 2018) involving 11,188 positive and negative PPIs derived from the *S. cerevisiae* core subset in the DIP database. We first validated the prediction performance of iPPI based on five-fold cross validation following You et al. (You, et al., 2014). Test results revealed comparable or even superior prediction performance on this dataset compared to the previous cross-validation results (Table SS1), which further validated that our model did not suffer from the overfitting problem. To further validate the generalization of our framework across different datasets and species, we additionally evaluated the prediction performance of iPPI on four additional datasets of other species, which was also derived from Zhou et al. (Zhou, et al., 2011) including *C. elegans* with 4,013 PPIs, *E.coli* with 6,954 PPIs, *H.* *sapiens* with 1,412 PPIs and *M. musculus* with 313 PPIs . We first trained a model based on 11,188 positive and negative PPIs from the above-mentioned *S. cerevisiae* core subset, which was constructed through our optimized deep learning framework (see the section of "Construction of predictors"), and the network parameters set by the hyperparameter calibration. Based on the results reported by DPPI (Hashemifar, et al., 2018), we found that the prediction performance of iPPI was comparable on other species datasets (Figure SS1), demonstrating the generalization capacity of our framework.

**References**

**Supplementary figures**

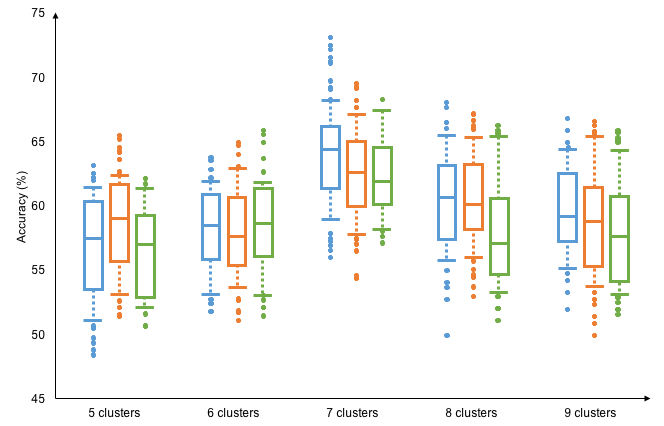

**Figure S1. The accuracy of prediction by traversing the cluster sizes from 5 to 9 and the sliding window sizes from 3 to 5 (represent by different colors from left to right). Each of cluster-available entries in AAindex was used to encode train dataset for each traversal.**

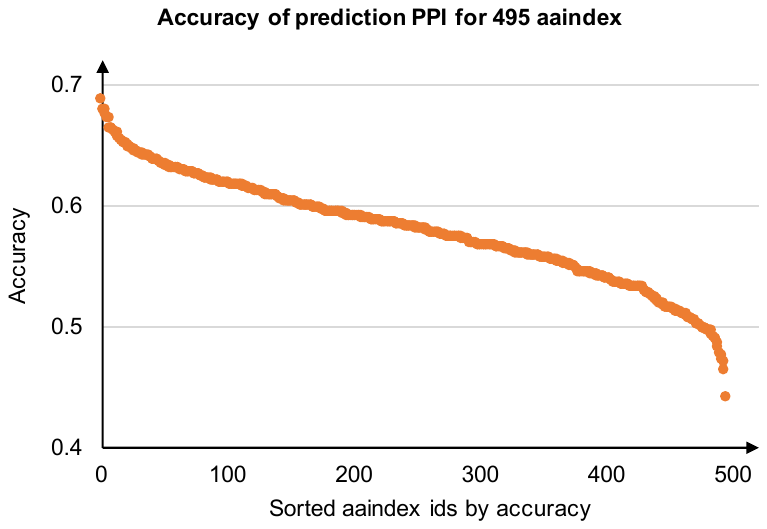

**Figure S2. The sorted accuracy of prediction by setting the window length as 3 and the cluster size as 7. By clustering properties, the train dataset encoded by each of 495 cluster-available entries in AAindex was used in this analysis.**

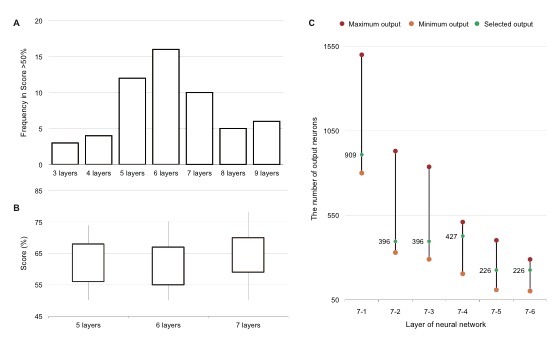

**Figure S3. The optimization processes of model for the size of network layers and the number of output neurons of each dense layer. (A) the distribution of the frequencies of model score >50% for the different size of network layers. (B) the scores (prediction accuracy) on the top three layer sizes of the frequencies. (C) the optimization of the number of output neurons for six dense layers of seven network layers. The interval of each dense layer was set from the optimization in the step (B). The hyperparameter calibration procedures were used in these optimization processes.**

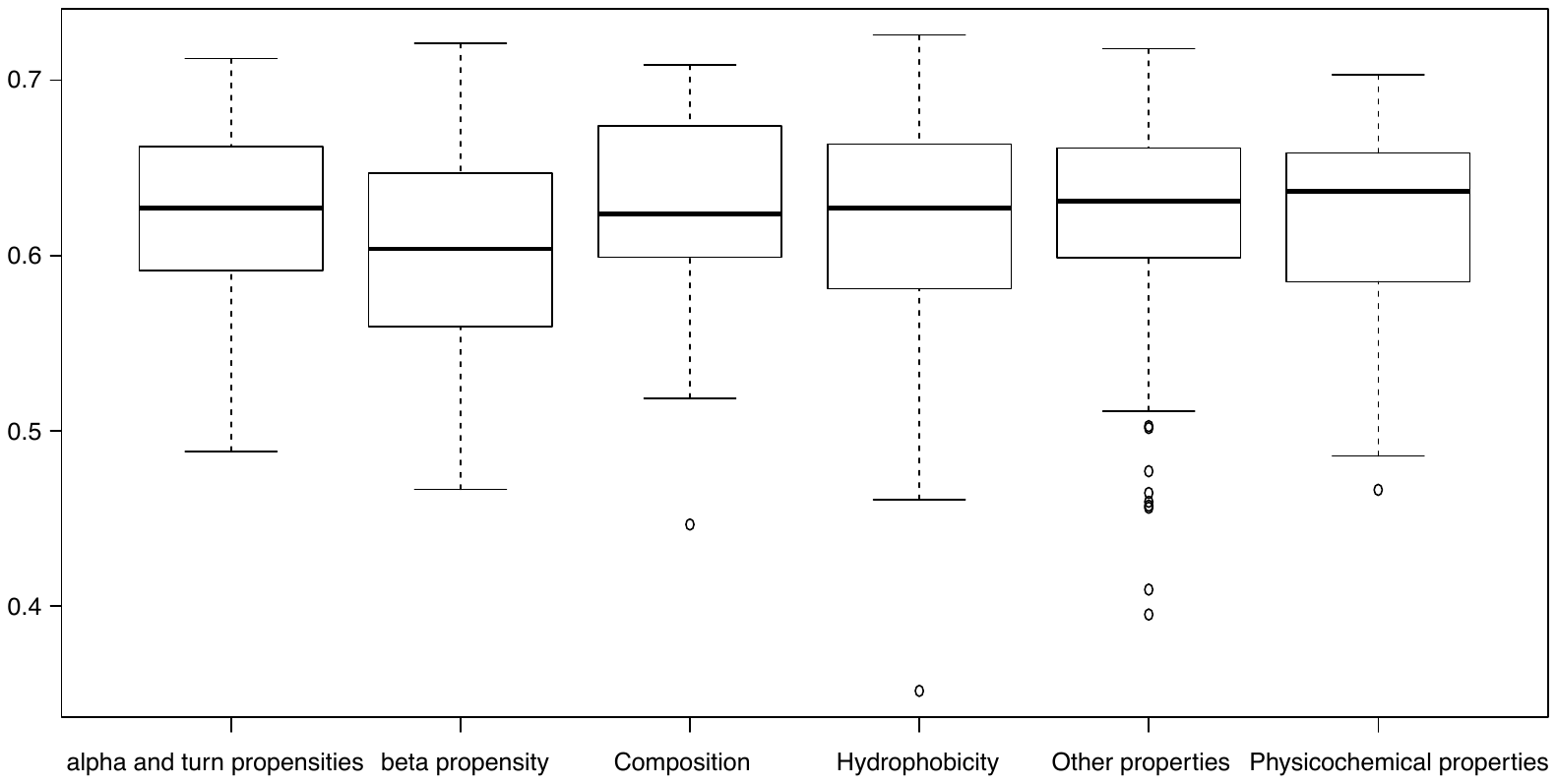

**Figure S4. The comparison of prediction capacities of the entries on different groups of property. The boxplot shows the predicted accuracy distribution of entries grouped in different categories. The prediction accuracy of all entries is derived from the results of Figure S1.**

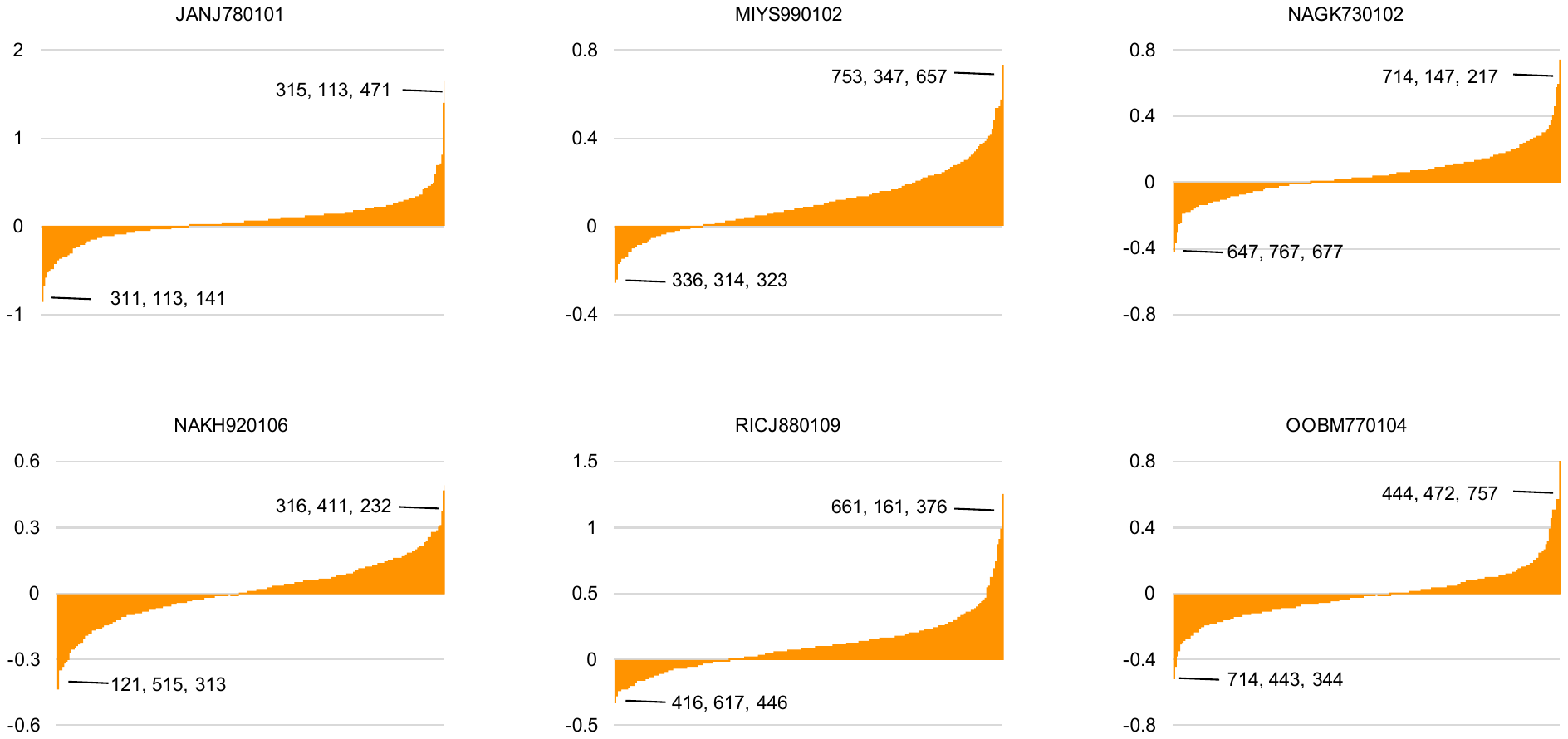

**Figure S5. The differences between positive and negative samples of CAM based on the encodings of the best 6 entries. The in vivo human PPI data from Hippie database v1.2 (Schaefer et al., 2012) mentioned in the main text was used to extract CAM. Each three-digit number in the figure represents a CAM, and each number in the three-digit number represents the cluster ID corresponding to entry. The enrichment value greater than zero indicates a positive sample, and vice versa.**

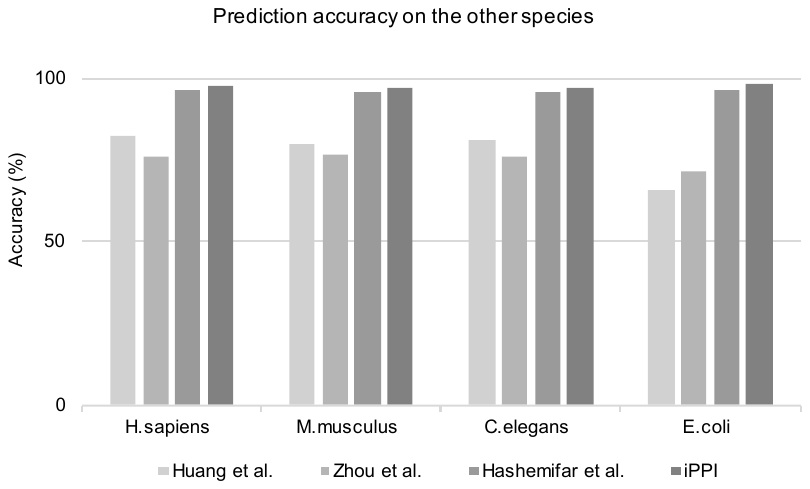

**Figure SS1. The comparison of prediction performance on other species datasets.**

**Supplementary tables**

**Table S1. The amino acids of each cluster for the top six entries of prediction accuracy in each group of AAindex. The sorted accuracy of prediction was obtained from Figure S2. The entries include** **JANJ780101 (Janin and Wodak, 1978),** **MIYS990102 (Miyazawa and Jernigan, 1999), NAGK730102 (Nagano, 1973),** **RICJ880109 (Richardson and Richardson, 1988),** **OOBM770104 (Oobatake and Ooi, 1977).**

| **AAINDEX ID** | **JANJ780101** | **MIYS990102** | **NAGK730102** | **NAKH920106** | **RICJ880109** | **OOBM770104** |
| --- | --- | --- | --- | --- | --- | --- |
| Cluster 1 | C | I, F, L | E | W, C | G, P | F, Y, W |
| Cluster 2 | V, A, I, G, L, F | C, W, Y, V, M | P, K, R, S, N | M, H, Y | S, E, Y, C | L, M, I, R, H |
| Cluster 3 | W, M | H, A | H, A, G, D | F, N, I | T, D, N, H | E, V |
| Cluster 4 | S, T | R, T, G | C, Y, W, Q | Q, T, D, P, V | K | Q |
| Cluster 5 | H, P, Y, N, D | P, S, N, Q | T, L, M | G, R, A | F, Q, L, R, I, V | D, T, N, P, K, C |
| Cluster 6 | E, Q | E, D | V, F | S, E | M, W | A, S |
| Cluster 7 | K, R | K | I | L, K | A | G |

**Table S2. The clustering values for the top six entries that associate with the clusters in Table S1.**

| **AAINDEX ID** | **JANJ780101** | **MIYS990102** | **NAGK730102** | **NAKH920106** | **RICJ880109** | **OOBM770104** |
| --- | --- | --- | --- | --- | --- | --- |
| CV 1 | 15.500 | -0.347 | 0.330 | 1.380 | 0.400 | -22.301 |
| CV 2 | 25.317 | -0.190 | 0.744 | 2.440 | 0.725 | -16.163 |
| CV 3 | 34.100 | 0.000 | 0.908 | 4.197 | 0.975 | -13.841 |
| CV 4 | 43.500 | 0.057 | 1.128 | 5.202 | 1.100 | -13.689 |
| CV 5 | 55.620 | 0.090 | 1.260 | 6.553 | 1.250 | -12.244 |
| CV 6 | 68.450 | 0.120 | 1.390 | 7.580 | 1.500 | -9.997 |
| CV 7 | 98.850 | 0.170 | 1.540 | 8.295 | 1.800 | -7.592 |

*Note:* CV denotes cluster value

**Table S3. the prediction results of each of selected entries in AAindex and their combination that implement by the hybrid deep neural network framework mentioned the manuscript (the parameters of the network were set by the hyperparameter calibration procedure with 1000 evaluations, respectively). These results were computed by 10-fold cross validations.**

| **Features** | **Accuracy** | **Precision** | **Recall** | **F1 Score** |
| --- | --- | --- | --- | --- |
| **JANJ780101** | 0.688±0.040 | 0.648±0.033 | 0.825±0.077 | 0.726±0.079 |
| **MIYS990102** | 0.679±0.018 | 0.640±0.049 | 0.817±0.086 | 0.718±0.008 |
| **NAGK730102** | 0.672±0.094 | 0.627±0.069 | 0.848±0.048 | 0.721±0.016 |
| **NAKH920106** | 0.657±0.079 | 0.616±0.053 | 0.835±0.031 | 0.709±0.078 |
| **RICJ880109** | 0.655±0.081 | 0.610±0.054 | 0.855±0.007 | 0.712±0.040 |
| **OOBM770104** | 0.672±0.094 | 0.642±0.002 | 0.777±0.034 | 0.703±0.022 |
| **Combined** | 0.747±0.036 | 0.764±0.015 | 0.871±0.083 | 0.809±0.081 |

**Table S4. The *p*-values were computed by the comparisons of prediction accuracy for selected six entries in AAindex and their combination. The data associate with Table S3.**

| **Features** | **JANJ780101** | **MIYS990102** | **NAGK730102** | **NAKH920106** | **RICJ880109** | **OOBM770104** | **Combined** |
| --- | --- | --- | --- | --- | --- | --- | --- |
| **JANJ780101** | 1.000 | 0.073 | 0.052 | 0.019 | 0.013 | 0.050 | 0.000 |
| **MIYS990102** |  | 1.000 | 0.138 | 0.036 | 0.036 | 0.131 | 0.000 |
| **NAGK730102** |  |  | 1.000 | 0.039 | 0.041 | 0.538 | 0.000 |
| **NAKH920106** |  |  |  | 1.000 | 0.693 | 0.041 | 0.000 |
| **RICJ880109** |  |  |  |  | 1.000 | 0.039 | 0.000 |
| **OOBM770104** |  |  |  |  |  | 1.000 | 0.000 |
| **Combined** |  |  |  |  |  |  | 1.000 |

**Table S5. The comparison of prediction capacities between CAM with six entries and CAM with 20 amino acids.**

| **Encoding** | **Accuracy** | **Precision** | **Recall** | **F1 Score** |
| --- | --- | --- | --- | --- |
| CAM with six entries (the combination)* | 0.747 | 0.764 | 0.871 | 0.809 |
| CAM with 20 amino acids | 0.531 | 0.524 | 0.567 | 0.545 |

**Table S6. The parameters of the deep learning framework of iPPI.**

| **Network terms** | **parameters** |
| --- | --- |
| Input features | 343*6 |
| learning rate | 1.4792765037709115E-5 |
| L2 | 1.978894473010271E-5 |
| SGD method | ADAM(Kingma and Ba, 2015) |
| N Layer 1 | 909 (0.9) |
| N Layer 2 | 396 (0.9) |
| N Layer 3 | 396 (0.9) |
| N Layer 4 | 427 (0.9) |
| N Layer 5 | 226 |
| N Layer 6 | 226 |

*Note:* SGD: Stochastic Gradient Descent; L2: L2 regularization constant; N: the number of output neurons (dropout rate)

**Table SS1. Performance comparison of iPPI with other state-of-the art methods on the *S. cerevisiae* core subset. Other tools reported results from (Hashemifar, et al., 2018). Note: Bold font is used to indicate the best performance.**

| Method | Precision (%) | Recall (%) | Accuracy (%) |
| --- | --- | --- | --- |
| Report by Yang (Yang, et al., 2010) | 90.24 | 81.03 | 86.15 |
| Report by Zhou (Zhou, et al., 2011) | 89.50 | 87.37 | 88.56 |
| Report by You (You, et al., 2013) | 87.59 | 86.15 | 87.00 |
| Report by You (You, et al., 2014) | 91.94 | 90.67 | 91.36 |
| Report by (Wong, et al., 2015) | 96.45 | 91.10 | 93.92 |
| Report by DPPI (Hashemifar, et al., 2018) | 96.68 | 92.24 | 94.55 |
| Report by iPPI | 94.95 | 93.41 | 94.72 |
